# Cardiac recovery from pressure overload is not altered by thyroid hormone status in old mice

**DOI:** 10.1101/2023.11.17.567523

**Authors:** Helena Kerp, Janina Gassen, Susanne Camilla Grund, Georg Sebastian Hönes, Stefanie Dörr, Jens Mittag, Nina Härting, Frank Kaiser, Lars Christian Moeller, Kristina Lorenz, Dagmar Führer

## Abstract

Thyroid hormones (TH) are known to have various effects on the cardiovascular system. However, the impact of TH levels on preexisting cardiac diseases are still unclear. Pressure overload due to arterial hypertension or aortic stenosis and aging are major risk factors for the development of structural and functional abnormalities and subsequent heart failure.

Here, we assessed the sensitivity to altered TH levels in aged mice with maladaptive cardiac hypertrophy and cardiac dysfunction induced by transverse aortic constriction (TAC). Mice at the age of 12 months underwent TAC and after induction of left ventricular pressure overload, received T4 or anti-thyroid medication in the drinking water over the course of 4 weeks.

T4 excess or deprivation in older mice had no or only very little impact on cardiac function (fractional shortening), cardiac remodeling (cardiac wall thickness, heart weight, cardiomyocyte size, apoptosis and interstitial fibrosis) and mortality. This is surprising, because T4 excess or deprivation had significantly changed the outcome after TAC in young 8-week-old mice.

In summary, our study shows that low and high TH availability have little impact on cardiac function and remodeling in older mice with preexisting pressure induced cardiac damage. This suggests that even though cardiovascular risk is increasing with age, the response to TH stress may be dampened in certain conditions.

## 2 Introduction

Despite great therapeutic advances, cardiovascular diseases remain the most common cause of death, globally (1). Many clinical and animal studies have described a close association of thyroid dysfunction with cardiovascular diseases (2-7). Both, systemic hyperthyroidism and hypothyroidism have been identified as risk factors for heart failure (8). Aging acts as an important further modifier, as both the prevalence for thyroid dysfunctions and cardiac diseases increases with age (9-11).

However, the relevance and interplay of altered thyroid hormone (TH) availability, age and their impact on progression or recovery of cardiac diseases are still not fully understood.

TH exert positive inotropic as well as chronotropic effects in the heart and reduce cardiac afterload by decreasing arterial vascular smooth muscle tone (3, 12-14). These TH effects involve the regulation of protein levels of sarcomeric proteins, such as myosin heavy chain alpha and beta, regulators of calcium homeostasis such as sarcoplasmic/endoplasmic reticulum Ca(2+)ATPase 2a2 and ryanodine receptor 2 (15-17). For a limited duration, treatment with TH has been shown to improve cardiac output, reduce afterload and leads to a compensated, so-called physiological or “athletic” cardiac hypertrophy (3, 18-24).

In our previous study in 8-week-old mice, we found a low TH state to be beneficial under conditions of left ventricular pressure overload induced by transverse aortic constriction (TAC). Reduced cardiac hypertrophy and improved cardiac function were evident after four weeks of TH deprivation. In contrast, TH excess was not beneficial in this situation as it did not prevent cardiac hypertrophy, increased apoptosis and had a negative impact on cardiac function (25).

Here, we aimed to investigate whether or to which extent older mice at the age of 12 months develop changes in heart function and morphology after TAC in response to TH deprivation or high dose TH availability. In line with Kerp *et al*. (25), modulation of thyroid status was started after one week of TAC when cardiac hypertrophy and first signs of cardiac depression have developed. In contrast to our previous study in young mice, low and high TH availability showed little impact on cardiac hypertrophy, apoptosis, fibrosis or cardiac function suggesting that even though cardiovascular risk is increasing with age, the response to TH stress may be dampened.

## 3 Material and Methods

### 3.1 Animals

Male wild-type C57BL/6JRj mice (Janvier Labs, France) were studied at the age of 12 months as previously described (25). Briefly, all mice were single-housed under standard conditions (room temperature 23 ±1°C; relative humidity 55±10%) and under a 12:12 hour light-dark cycle. Standard chow diet and water were provided *ad libitum*. All animal experiments were performed in accordance with the German regulations for Laboratory Animal Science (GV-SOLAS) and the European Health Law of the Federation of Laboratory Animal Science Associations (FELASA). The experimental protocols were approved by the Landesamt für Natur, Umwelt und Verbraucherschutz Nordrhein-Westfalen, Germany (LANUV-NRW, AZ 84-02.04.2016.A261).

### 3.2 Transverse aortic constriction

Mice were subjected to TAC using a 25-gauge needle to induce chronic left ventricular pressure overload as previously described (26, 27). One day prior to TAC and 1, 3, and 5 weeks after TAC echocardiography was performed (Fig. 1A). Mice with an aortic pressure gradient below 60 mmHg at 1 week after TAC were excluded from the study.

**Figure 1:**
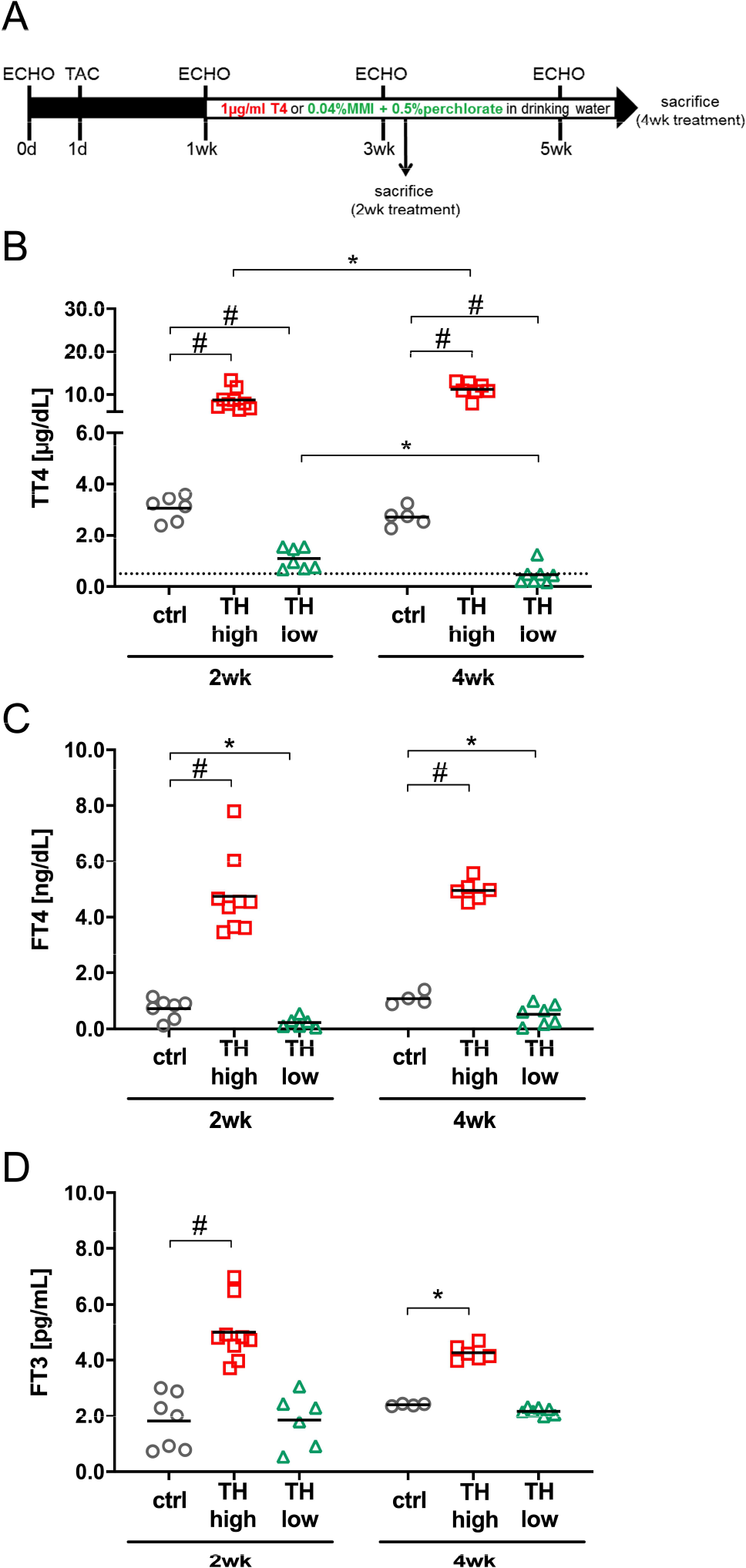
Study design and serum TH status. 12-month-old male C57Bl/6 mice were subjected to transaortic constriction (TAC) at experimental day 1. Echocardiographic measurements (ECHO) were conducted one day prior to TAC (day 0) and after 1, 3 and 5 weeks. One week after TAC, oral T4 treatment (TH high) and TH deprivation (MMI/ClO_4_^-^ in combination with low iodine diet [TH low]) was started. Controls (ctrl) received drinking water without supplements. Mice were sacrificed 2 or 4 weeks after induction of TH dysfunction (**A**). TT4 (**B**), FT4 (**C**) and FT3 (**D**) serum concentrations confirmed TH excess and TH deprivation at both time points in the two treatment groups compared to controls (**C**). Dotted line represents detection limit of assay and values below were calculated from the standard curve. \**p*<0.05, ^#^*p*<0.0001 by Two-Way ANOVA and Tukey’s *post hoc* analysis, ctrl=control, wk=weeks.

### 3.3 Echocardiography

Transthoracic echocardiograms were performed in a blinded fashion using the Vevo3100 highresolution imaging system (VisualSonics) and a 30-MHz probe with pentobarbital (35mg/kg body weight, i.p.) as anaesthetic. 2D M-mode images in the short axis view at the proximal level of the papillary muscles were used to evaluate end-diastolic and -systolic intraventricular septal (IVSd/IVSs), left ventricular posterior wall thickness (LVPWd/LVPWs) and end-diastolic and - systolic left ventricular internal diameters (LVIDd/LVIDs). Pulsed-wave doppler measurements were performed to evaluate the peak blood flow velocities at the site of constriction (V_max_ (mm/sec)).

Fractional shortening (FS) and aortic pressure gradients (mmHg) were calculated using the VisualSonics Cardiac Measurements software. The data represent the average of at least six cardiac cycles. The investigators were blinded regarding treatment groups during measurements and data analysis. Measurements at heart rates below 450 bpm were excluded.

### 3.4 Treatment

Mice were subjected to two different treatment protocols: A) To increase TH availability (TH high) mice received thyroxine (T4) in drinking water (1 μg/ml, Sigma-Aldrich (T2376), USA; stock solution: 100 µg/ml T4 solved in 40 mM NaOH and 2 g/l bovine serum albumin). B) For TH deprivation (TH low), mice received a low-iodine diet (LoI; MD.1571, Envigo, USA) and drinking water supplemented with 0.04% methimazole (MMI, Sigma-Aldrich (301507), USA), 0.5% sodium perchlorate (ClO_4_^-^, Sigma-Aldrich (310514), USA) and 0.3% saccharine as sweetener (Sigma-Aldrich (240931), USA) (LoI/MMI/ClO_4_^-^). In the control group (ctrl), animals were fed a normal diet (MD.1572, Envigo, USA) and received drinking water without supplements.

Except for the low iodine content in the TH deprivation group, caloric and nutritional composition of the diet was comparable for all mice (MD.1572, Envigo, USA, for control and T4 treated mice). The indicated treatment started one week after TAC surgery and was continued for two or four weeks (Fig. 1A).

### 3.5 Organ isolation and serum measurements

For organ extraction, mice were anaesthetized by i.p. injection of 200 µl Ketamine/Xylazine mixture (150 µl of 100 mg/ml Ketamine (Beta-pharm, Germany) and 50 µl of 20 mg/ml Xylazine (Ceva, Germany) and blood was withdrawn by cardiac puncture. Mice were perfused with heparinized saline by transcardial perfusion. Hearts were shock frozen in liquid N_2_ and stored at −80 °C or fixed in 4% buffered formalin (Formafix, Germany). Blood samples were placed on ice for 30 min, centrifuged and concentrations of free triiodothyronine (FT3), free T4 (FT4) and total T4 (TT4) in mouse sera were measured according to the manufacturer’s instructions of commercial ELISA kits (DRG Instruments GmbH, Marburg, Germany) as previously described (25, 28, 29). Detection limit was 0.5 µg/dL, 0.05 ng/dL and 0.05 pg/mL for TT4, FT4 and FT3, respectively. Values were calculated from the standard curve.

### 3.6 RNA isolation and qRT-PCR

Total cardiac RNA was isolated and reverse transcribed to cDNA as previously described (29). In compliance with the guidelines for RT-PCR of MIQE, we used three reference genes to assure correct normalization and calculation, *Gapdh* (glyceraldehyde-3-phosphate dehydrogenase), *Rn18s* (18S ribosomal RNA) and *Polr2a* (polymerase RNA II). Primer sequences are listed in Supplementary Table 1. Analysis and calculation of the fold changes of gene expression were applied to Ct-values ≤35.

### 3.7 Histological staining

Formalin fixed hearts were embedded in paraffin. 5 µm sections were used for staining. Hematoxylin and eosin (H&E) staining was used for determination of cardiomyocyte size and Sirius Red for the analysis of fibrosis as described previously (26). For quantification, stained sections were imaged using an Olympus BX51 microscope (Olympus Life Science, Tokyo, Japan). Cross-sectional areas of cardiomyocytes were determined (n= 37-88 cells per animal) using ImageJ. Only cells with central nucleus were included in the analysis. Fibrosis was calculated as ratio of red stained/myocardial area via Adobe Photoshop. Analysis and quantification were performed by blinded researchers.

### 3.8 TUNEL assay

After dewaxing and rehydration, cardiac sections were permeabilized using proteinase K. For TUNEL (terminal deoxynucleotidyl transferase dUTP nick-end labeling) staining, an *in situ* detection kit was used according to the manufacturer’s protocol (Sigma-Aldrich, USA). Sample pretreatment with DNase I served as positive control and TUNEL reaction mixture without terminal transferase (TdT) as negative control. DAPI (D1306, Thermo Fisher Scientific, USA) and wheat germ agglutinin (W11261, Thermo Fisher Scientific, USA) were used as counterstains for cell nuclei and membranes. Olympus BX51 upright microscope was used for the imaging and quantification was conducted in a blinded manner.

### 3.9 Statistical analysis

GraphPad Prism 6 Software was used for statistical analysis. Two-Way ANOVA followed Tukey’s *post hoc* analysis was used. P-values of \**p*<0.05, \*\**p*<0.01, \*\*\**p*<0.001, _#_*p*<0.0001 were considered significant. Outliers were identified using Graph Pad Outlier test (excluded if *p*<0.05).

## 4 Results

TH availability may alter the progression of heart disease and the development of heart failure (2, 5, 30). To investigate which impact aging has on this process, we subjected 12-month-old male mice to chronic left ventricular pressure overload induced by ligation of the transverse aorta. Well-characterized features of this model are pathological growth of the heart, i.e. maladaptive cardiac hypertrophy associated with interstitial fibrosis and apoptosis. Subsequently fractional shortening (FS) decreases and congestive heart failure develops (31).

Changes in TH availability were induced by T4 administration or anti-thyroidal drug treatment (LoI/MMI/ClO_4_-), respectively, initiated 1 week after TAC surgery and continued over 4 weeks as depicted in the study design (Fig. 1A). As expected, increased and decreased TH serum concentrations were measured in the mice reflecting states of TH excess and TH deprivation at 2 and 4 weeks after start of treatment (Fig. 1B-D).

### 4.1 TH effects on maladaptive cardiac hypertrophy in adult mice

Serial echocardiographic measurements were performed to monitor the progression of cardiac hypertrophy and cardiac dysfunction in response to TAC. Aortic pressure gradients induced by TAC were comparable in the different treatment groups and were stable throughout the experiment (Fig. 2A). At 1 week of TAC and before manipulation of TH status commenced, a significantly decreased FS was confirmed. Of note, the subsequent alteration of TH levels, i.e. gradual TH increase and TH deprivation, did not alter FS over 4 weeks of treatment (Fig. 2B, Table 1). Thickness of left ventricular wall was significantly increased after 1 week of TAC as depicted by both posterior wall and septum diameters compared to baseline values and hardly changed during the following 4 weeks of observation. Importantly, this course of wall thickness development was independent of changes in serum TH levels as it was comparable for all treatment and control groups. A slight, nonsignificant decrease in wall thickness was observed after 4 weeks of TH deprivation (Fig. 2C, D), however, the left ventricular inner diameter was unaltered at all time points and in all treatment and control groups (Suppl. Fig. 1A).

**Figure 2:**
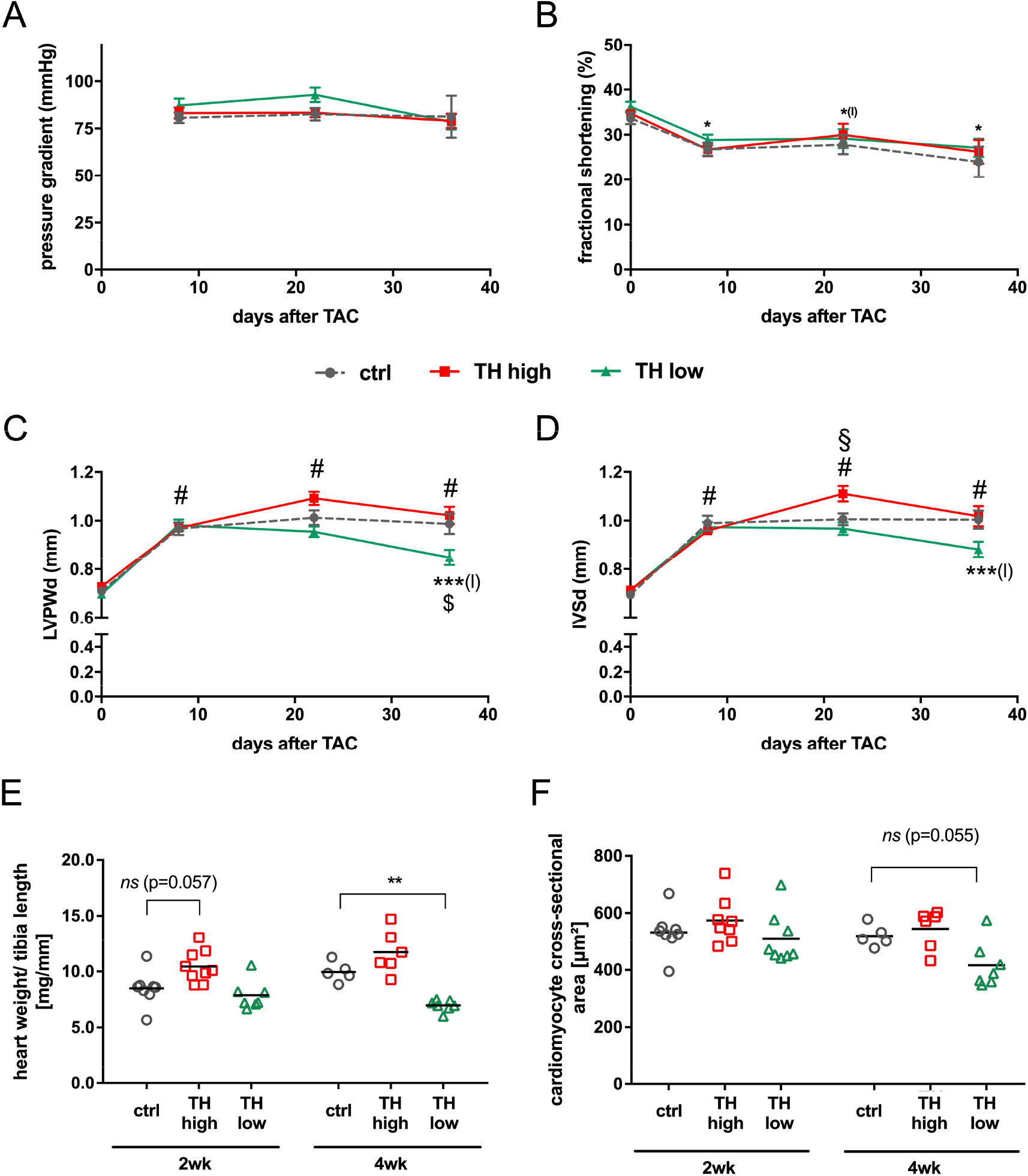
Cardiac function and hypertrophy characteristics. Echocardiography was conducted prior TAC and 1, 3 and 5 weeks after surgery. After 1 week, pressure gradients confirmed successful TAC surgery (**A**) and fractional shortening decreased in all indicated groups compared to baseline values without an apparent impact of thyroid status (**B**). For cardiac hypertrophy analysis, diastolic left ventricular posterior wall (LVPWd; **C**) and interventricular septum (IVSd; **D**) thickness were calculated, in addition to heart weight to tibia length ratio (**E**) and cardiomyocyte cross-sectional area (**F**). TH deprivation reduced cardiac hypertrophy parameters (LVPWD, IVSd and heart weight to tibia length). Values are indicated as mean ± SEM, for n see Supplemental Table 1. *^(l)^*p*<0.05 (TH low vs basal),\**p*<0.05, \*\**p*<0.01, ***^(l)^*p*<0.001 (TH low vs basal), ^#^*p*<0.0001 (for echocardiographic parameters all groups vs basal) by Two-Way ANOVA and Tukey’s *post hoc* analysis, ctrl=control, wk=weeks., ***^(l)^*p*<0.001 (TH low vs basal), ^#^*p*<0.0001 (for echocardiographic parameters all groups vs basal), $*p*<0.05 (TH low vs ctrl at 5 weeks after TAC) and §*p*<0.05 (TH high vs ctrl at 3 weeks after TAC) by Two-Way ANOVA and Tukey’s *post hoc* analysis, ctrl=control, wk=weeks.

**Table 1:** Reduction of FS over time in 12 months compared to 8-week-old groups (23), receiving the same experimental conditions of TAC surgery and modulation of TH status (TH high: T4 treatment and TH low: TH deprivation).

|  | 8 weeks old | 12 months old |
| --- | --- | --- |
| 1 week after TAC vs basal | 20.7 ± 2.40% | 18.8 ± 2.76% |
| 5 weeks after TAC vs basal<br>(control) | 33.9 ± 4.69% | 33.4 ± 10.27% |
| 5 weeks after TAC vs basal<br>(TH high) | 46.2 ± 3.51% | 27.8 ± 6.82% |
| 5 weeks after TAC vs basal<br>(TH low) | 19.7 ± 3.65% | 28.0 ± 6.72% |

These echocardiographic data on cardiac hypertrophy were validated by heart weight to tibia length ratios and cardiomyocyte cross-sectional area. Here a significantly reduced heart weight to tibia length ratio was found under TH deprivation (Fig. 2E) and a similar trend was observed for the cardiomyocyte cross-sectional area (Fig. 2F). No significant differences were detected after 2 and 4 weeks of T4 treatment (Fig. 2E, F). Thus, elevated TH serum concentrations did not affect the development of cardiac hypertrophy, whereas low TH serum concentrations had a mild impact on progression of cardiac hypertrophy. Neither cardiac function nor lung weight as a measure of cardiac congestion were influenced by altered TH levels during the 4-week treatment interval after induction of left ventricular pressure overload (Suppl. Fig. 1B).

### 4.2 TH-dependent development of fibrosis and apoptosis after TAC in 12-month-old mice

Apart from cardiac hypertrophy, deposition of collagen contributing to fibrosis and cardiomyocyte apoptosis are key features of pathological cardiac remodeling, which leads to cardiac dysfunction, altered electrophysiological dynamics and cardiac tissue stiffness (32). Interstitial fibrosis was histologically analyzed by Sirius Red staining and apoptotic cell death by TUNEL staining (27). No significant alterations in the extent of fibrosis (Fig. 3A, B) and apoptosis (Fig. 3C) were evident between the treatment groups after two and four weeks. Furthermore, cardiac expression of atrial natriuretic peptide (*Anp*) and brain natriuretic peptide (*Bnp*), markers that correlate with the development of heart failure and pathological cardiac hypertrophy, were analyzed by qRT-PCR (33, 34). *Anp* expression levels did not change in response to altered TH status, while cardiac *Bnp* expression was significantly lowered after 4 weeks of TH deprivation compared to controls (Fig. 3D, E).

**Figure 3:**
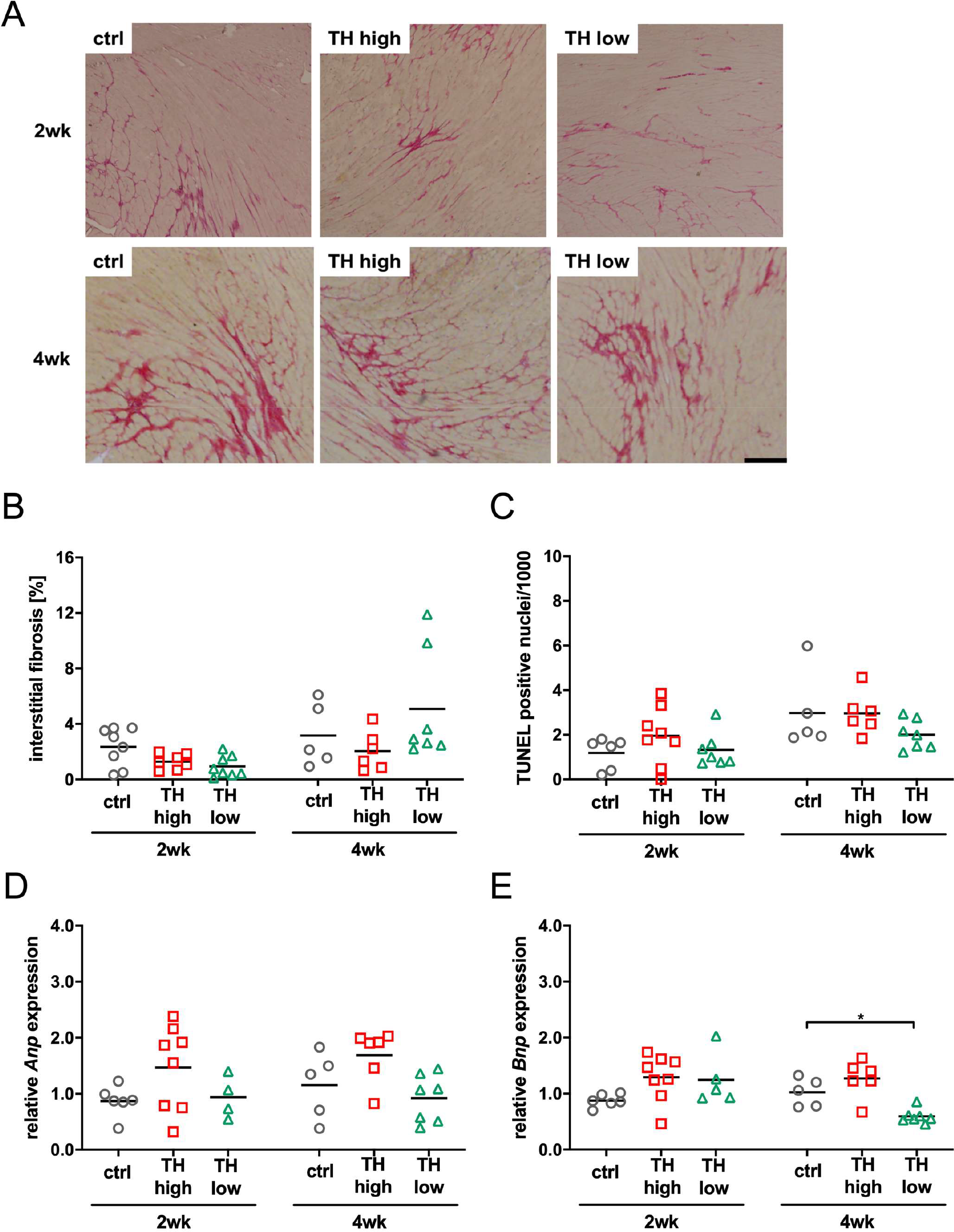
Analysis of fibrosis, apoptosis and cardiac stress markers. Representative pictures of Sirius Red stained sections (**A**, scale bar=200 µm) and relative quantification of interstitial fibrosis (**B**) are shown. TUNEL positive nuclei were counted to assess apoptotic rate (**C**) and *Anp* and *Bnp* expression as cardiac stress markers (**D, E**). TH deprivation reduced cardiac *Bnp* expression in 12-month-old mouse hearts at 5 weeks after TAC. Scatter dot plot and mean in all panels, \**p*<0.05 by Two-Way ANOVA and Tukey’s *post hoc* analysis, ctrl=control, wk=weeks.

Moreover, despite the absence of apparent features of heart failure, three death events occurred in the controls and two in the T4 treatment group in comparison to none in TH deprived mice within 4 weeks of modulation of TH status.

### 4.3 TH responsiveness of TAC subjected hearts

To answer whether the higher or lower TH availability in the mice translated into altered expression of genes associated with hypo-or hyperthyroidism (13), TH target gene expression of myosin heavy chain alpha (*Myh6*) and beta (*Myh7*), ATPase sarcoplasmic/endoplasmic reticulum Ca2+ transporting 2 (*Serca2a2*) and ryanodine receptor 2 (*Ryr2*) was analyzed in mouse hearts, sacrificed after 2 and 4 weeks of treatment. As observed upon TH deprivation in previous studies (25), *Myh6* levels were significantly lowered and *Myh7* expression increased after 2 and 4 weeks of low TH treatment. High TH treatment did not affect *Myh6* expression but resulted in reduced expression of *Myh7* (Fig. 4A, B). Expression of *Ryr2* was decreased in the low TH group only after 2 weeks of treatment, whereas expression of *Serca2a2* was significantly reduced after 2 and 4 weeks under TH deprivation (Fig. 4B, C). Thus, both treatments leading to high or low TH serum concentration, respectively, caused the expected gene alteration in cardiac muscle. However, the effect on cardiac TH target gene expression was greater under TH deprivation than under T4 treatment. Expression analysis of *Thra* and *Thrb* encoding for TH receptor alpha (TRα) and beta (TRβ), respectively, confirmed the predominant expression of TRα over TRβ in heart (Suppl. Fig. 2). Expression of TRα was decreased under high TH and increased under low TH while TRβ was unaffected. Interestingly, an analysis of TRα isoform expression revealed that the proportion of TRα2, the isoform that is not able to bind TH, increased after 4 weeks of treatment, irrespective of the treatment regime (Fig. 4E, F).

**Figure 4:**
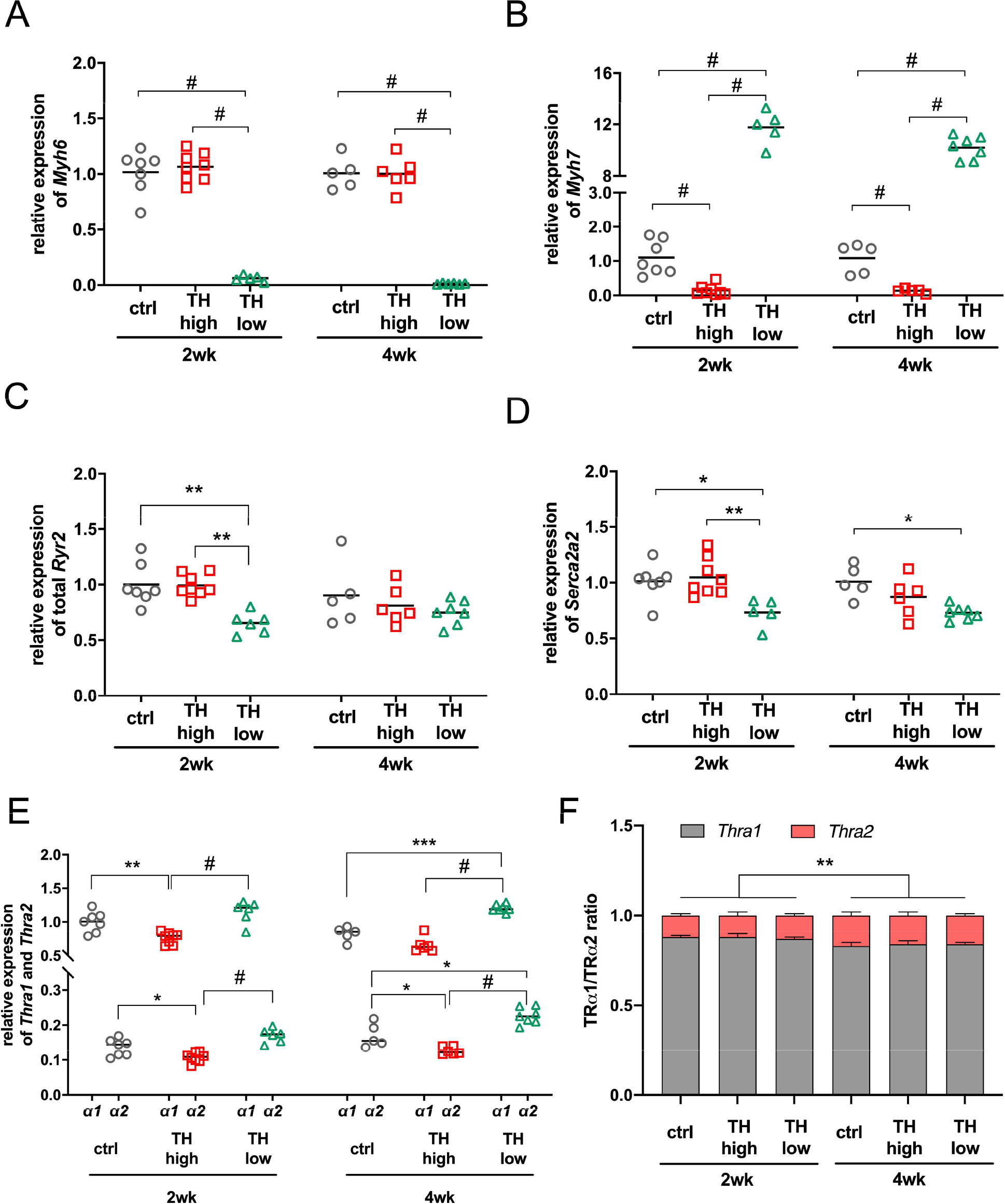
Expression of TH responsive genes in hearts of mice subjected to TAC and subsequent modulation of TH status by T4 treatment (TH high) or TH deprivation (TH low). Amount of *Myh6* (**A**), *Myh7* (**B**), *Ryr2* (**C**), *Atp2a2* (**D**), *Thra1* and *Thra2* (**E**) transcripts were determined in mouse hearts by qRT-PCR after 2 and 4 weeks of treatment. Expression ratio of TRα isoform 1 and 2 (**F**). Scatter dot plot and mean in all panels, One-Way ANOVA with Tukey’s *post-hoc* test, \**p*<0.05, \*\**p*<0.01, \*\*\**p*<0.01, ^#^*p*<0.0001, ctrl=control, wk=weeks.

## 5 Discussion

In our study, we investigated the impact of high and low TH availability after pressure overload induced cardiac dysfunction in male 12-month-old mice. The aim of the study was to delineate the sensitivity to altered TH levels in aged mice in the context of “pre-damaged” hearts with left ventricular pressure overload.

Chronic pressure overload is a pathological condition that triggers the development of heart failure. Left ventricular pressure overload can be caused by arterial hypertension or aortic stenosis and TAC as a mouse model mimicking such a condition is widely used. TAC induces the development of cardiac hypertrophy, fibrosis, an increase in apoptosis and eventually cardiac dilatation and congestive heart failure, characteristic features of heart disease induced by chronic pressure overload (35).

Previously, we could show that modulation of the TH status one week after TAC surgery in young, 8-week-old mice impressively impacted the outcome of cardiac recovery. In contrast to controls, TH deprivation in young mice stopped the progression of cardiac hypertrophy and cardiac dysfunction with mild improvement of FS, whereas elevated TH availability increased apoptosis, did not prevent hypertrophy and decreased FS (25).

These results prompted us to add aging as another important modifier to this experimental setup since the prevalence of cardiac and TH related diseases increases with age (10, 11) and sensitivity to TH and cardiovascular outcome change with ageing. TH action in aged mice has been well documented and findings from our previous studies suggested decreased systemic effects in aged animals (28, 29). The heart as one of the main TH target organs was more likely to have a preserved TH action compared to other organs (29, 36). However, chronic TH excess in old mice led to a similar extent of cardiac hypertrophy, tachycardia and regulation of TH responsive gene expression as in young mice, whereas chronic TH deprivation resulted in an attenuated cardiac phenotype with less pronounced bradycardia and degree of gene regulation (28).

Effects of aging on cardiac hypertrophy and cardiac disease in animal models have been occasionally reported. Hypertrophic response of the left ventricle 4 weeks after ascending aortic banding in rats of 9 and 18 months of age showed a diminished degree of myocardial hypertrophy in old rats in terms of muscle weight, size of myocytes and protein content (37-39). Mouse studies reported contradictory results of either worse response to chronic pressure overload and lower survival of old animals (40) or less left ventricular remodeling and lower mortality than young mice after TAC (41). However, so far the impact of TH on TAC induced heart disease has not been studied in the context of ageing.

Due to the limited availability of aged TAC mouse models, our first aim was to identify similar conditions in 12-month-old as compared to 8-week-old mice for our setup so that aortic pressure gradients would not differ in older mice. This was achieved by using a bigger needle size (25G) during TAC, which resulted in a similar decline in FS within one week after surgery in aged compared to our previous study in young mice (25) and confirmed our approach to achieve age-independent conditions for TAC within the experimental time prior to TH alterations. 4 weeks of T4 or LoI/MMI/ClO_4_^-^ treatment resulted in different effects in both age groups, with worsening in young TH high and improvement in young TH low groups but little impact in old mice (Table 1).

Thus, TH alterations had no obvious detrimental or beneficial effects within the studied time frame in older mice. As no age effect was noted in the control group, we consider these findings unlikely a result of inappropriate experimental setting.

Another influencing aspect might be that treatment time and dosage has to be age-adjusted to achieve comparable effects in younger and older individuals. Of note, the modulation of TH status in the present study led to similar changes in TH serum concentrations and cardiac gene expression like observed in young mice, further supporting comparable experimental conditions between the study in young and the herein reported study in older mice (25).

Early studies described a delayed development of cardiac hypertrophy in senescent mice upon thyroxine treatment. However, already after 9 days of treatment, no age-differences were apparent anymore. In addition, different dosages were tested and even with the lowest (1.7 µg/g BW) no age dependency of cardiac hypertrophy was noted (42). TH deprivation has been studied less frequently. Thus, it remains speculative whether treatment time or dosage would affect the outcome of TAC in older mice. In addition, other studies suggested that cardiac hypertrophy significantly depends on indirect TH action in the CNS rather than direct TH effects on heart tissue (43, 44). Thus, the observed age difference could also arise from altered central TH rather than cardiac TH signaling in young vs. old mice with induced pressure overload (45). Along the same lines, it is known that aging per se alters TH dependent gene expression in heart tissue (35). In addition, a higher responsiveness of heart rate was previously observed in hyperthyroid old compared to young mice.

Most clinical studies include patients with differing ages, co-morbidities, and different causes of cardiac dysfunction as well as common endpoints as all-cause mortality or cardiovascular death in correlation with the thyroid status so that conclusions for therapeutic TH modulation are difficult (46-48). TH modulation may well be of different impact in different subgroups of heart failure patients, e.g. in heart failure patients with or without atrial fibrillation (47) and in patients with congestive or dilative heart failure. Noteworthy, we detected a shift in the TRα-isoform ratio towards more TRα2 expression. TRα2 is incapable of binding TH and thus has a dominant-negative effect on TRα1 action (49). This increase was not affected by TH modulation, i.e. TH deprivation and TH treatment. However, as there was more TRα2 in hearts of all groups of four weeks of treatment, we suggest that the amount of TRα2 correlates with the hypertrophy progression. This is supported by previous studies that showed, a shift from TRα1 to TRα2 expression in the human heart during heart failure (50, 51). Moreover, elevated TRα2 expression was found to attenuate TRα1-mediated hypertrophy in this condition (52). These findings suggest that the TRα1:TRα2 ration might contribute to the outcome of TH modulation in the pathophysiology of heart disease.

The data of our current mouse study are focused and limited to TH sensitivity in 12-month-old mice after pressure overload induced hypertrophy and cardiac dysfunction but suggest that TH modulation (high and low levels) at older age – at least within a certain time frame – has less impact to foster key risk factors of heart failure.

Taken together, certain subgroups of patients with preexisting cardiovascular disease may respond to manipulation of TH status with a beneficial or detrimental outcome. Our study suggests different outcome to TH modulation based on age, thus the recognition of differences due to age, preexisting cardiovascular diseases and risk factors needs to be included in future studies.

## 7 Author Contributions

DF and KL planned the study and HK, JG, GSH and KL conducted the experiments. SCG, SD and NH assisted in data analysis and JM, LCM and FK contributed to interpretation of data. HK, DF and KL wrote the first manuscript draft. All authors contributed with further suggestions for presentation of results and discussion and approved the final version.

## 9 Acknowledgments

Authors thank the Imaging Core Facility Essen (IMCES) for expert technical assistance. We are also grateful to K. Schättel, S. Rehn and A. Jaeger for their dedicated technical support. We acknowledge support by the Open Access Publication Fund of the University of Duisburg-Essen.

## 11 Data Availability Statement

All relevant data is contained within the article. The original contributions presented in the study are included in the article/supplementary material. Further inquiries can be directed to the corresponding author/s.

## References

1. Bragazzi NL, Zhong W, Shu J, Abu Much A, Lotan D, Grupper A, et al. Burden of Heart Failure and Underlying Causes in 195 Countries and Territories from 1990 to 2017. Eur J Prev Cardiol (2021) 28(15):1682–90. Epub 2021/02/12. doi: 10.1093/eurjpc/zwaa147.

2. Dillmann W. Cardiac Hypertrophy and Thyroid Hormone Signaling. Heart failure reviews (2010) 15(2):125–32. Epub 2009/01/07. doi: 10.1007/s10741-008-9125-7.

3. Pantos C, Mourouzis I, Cokkinos DV. New Insights into the Role of Thyroid Hormone in Cardiac Remodeling: Time to Reconsider? Heart Fail Rev (2011) 16(1):79–96. Epub 2010/07/30. doi: 10.1007/s10741-010-9185-3.

4. Jabbar A, Pingitore A, Pearce SH, Zaman A, Iervasi G, Razvi S. Thyroid Hormones and Cardiovascular Disease. Nat Rev Cardiol (2017) 14(1):39–55. Epub 2016/11/05. doi: 10.1038/nrcardio.2016.174.

5. Razvi S, Jabbar A, Pingitore A, Danzi S, Biondi B, Klein I, et al. Thyroid Hormones and Cardiovascular Function and Diseases. J Am Coll Cardiol (2018) 71(16):1781–96. Epub 2018/04/21. doi: 10.1016/j.jacc.2018.02.045.

6. Debmalya S, Saumitra R, Singh MH. Interplay between Cardiovascular and Thyroid Dysfunctions: A Review of Clinical Implications and Management Strategies. Endocr Regul (2022) 56(4):311–28. Epub 2022/10/22. doi: 10.2478/enr-2022-0033.

7. Paschou SA, Bletsa E, Stampouloglou PK, Tsigkou V, Valatsou A, Stefanaki K, et al. Thyroid Disorders and Cardiovascular Manifestations: An Update. Endocrine (2022) 75(3):672–83. Epub 2022/01/16. doi: 10.1007/s12020-022-02982-4.

8. Cappola AR, Desai AS, Medici M, Cooper LS, Egan D, Sopko G, et al. Thyroid and Cardiovascular Disease: Research Agenda for Enhancing Knowledge, Prevention, and Treatment. Circulation (2019) 139(25):2892–909. Epub 20190513. doi: 10.1161/CIRCULATIONAHA.118.036859.

9. Barbesino G. Thyroid Function Changes in the Elderly and Their Relationship to Cardiovascular Health: A Mini-Review. Gerontology (2019) 65(1):1–8. Epub 2018/07/23. doi: 10.1159/000490911.

10. Waring AC, Arnold AM, Newman AB, Buzkova P, Hirsch C, Cappola AR. Longitudinal Changes in Thyroid Function in the Oldest Old and Survival: The Cardiovascular Health Study All-Stars Study. J Clin Endocrinol Metab (2012) 97(11):3944–50. Epub 2012/08/11. doi: 10.1210/jc.2012-2481.

11. Writing Group M, Mozaffarian D, Benjamin EJ, Go AS, Arnett DK, Blaha MJ, et al. Executive Summary: Heart Disease and Stroke Statistics--2016 Update: A Report from the American Heart Association. Circulation (2016) 133(4):447–54. Epub 2016/01/27. doi: 10.1161/CIR.0000000000000366.

12. Li M, Iismaa SE, Naqvi N, Nicks A, Husain A, Graham RM. Thyroid Hormone Action in Postnatal Heart Development. Stem Cell Res (2014) 13(3 Pt B):582–91. Epub 2014/08/05. doi: 10.1016/j.scr.2014.07.001.

13. Dillmann WH. Biochemical Basis of Thyroid Hormone Action in the Heart. Am J Med (1990) 88(6):626–30. Epub 1990/06/01. doi: 10.1016/0002-9343(90)90530-q.

14. Kahaly GJ, Dillmann WH. Thyroid Hormone Action in the Heart. Endocr Rev (2005) 26(5):704–28. Epub 2005/01/06. doi: 10.1210/er.2003-0033.

15. Holt E, Sjaastad I, Lunde PK, Christensen G, Sejersted OM. Thyroid Hormone Control of Contraction and the Ca(2+)-Atpase/Phospholamban Complex in Adult Rat Ventricular Myocytes. J Mol Cell Cardiol (1999) 31(3):645–56. Epub 1999/04/13. doi: 10.1006/jmcc.1998.0900.

16. Kiss E, Jakab G, Kranias EG, Edes I. Thyroid Hormone-Induced Alterations in Phospholamban Protein Expression. Regulatory Effects on Sarcoplasmic Reticulum Ca2+ Transport and Myocardial Relaxation. Circ Res (1994) 75(2):245–51. Epub 1994/08/01.

17. Dore R, Watson L, Hollidge S, Krause C, Sentis SC, Oelkrug R, et al. Resistance to Thyroid Hormone Induced Tachycardia in Rthalpha Syndrome. Nat Commun (2023) 14(1):3312. Epub 20230607. doi: 10.1038/s41467-023-38960-1.

18. Shimizu I, Minamino T. Physiological and Pathological Cardiac Hypertrophy. Journal of molecular and cellular cardiology (2016) 97:245–62. Epub 2016/06/06. doi: 10.1016/j.yjmcc.2016.06.001.

19. Klein I, Danzi S. Thyroid Disease and the Heart. Circulation (2007) 116(15):1725–35. Epub 2007/10/10. doi: 10.1161/CIRCULATIONAHA.106.678326.

20. Thomas TA, Kuzman JA, Anderson BE, Andersen SM, Schlenker EH, Holder MS, Gerdes AM. Thyroid Hormones Induce Unique and Potentially Beneficial Changes in Cardiac Myocyte Shape in Hypertensive Rats near Heart Failure. Am J Physiol Heart Circ Physiol (2005) 288(5):H2118–22. Epub 2004/12/18. doi: 10.1152/ajpheart.01000.2004.

21. Gerdes AM, Iervasi G. Thyroid Replacement Therapy and Heart Failure. Circulation (2010) 122(4):385–93. Epub 2010/07/28. doi: 10.1161/circulationaha.109.917922.

22. Gerdes AM, Moore JA, Hines JM. Regional Changes in Myocyte Size and Number in Propranolol-Treated Hyperthyroid Rats. Lab Invest (1987) 57(6):708–13. Epub 1987/12/01.

23. Ito K, Kagaya Y, Shimokawa H. Thyroid Hormone and Chronically Unloaded Hearts. Vascul Pharmacol (2010) 52(3-4):138–41. Epub 2009/11/03. doi: 10.1016/j.vph.2009.10.004.

24. Yao J, Eghbali M. Decreased Collagen Gene Expression and Absence of Fibrosis in Thyroid Hormone-Induced Myocardial Hypertrophy. Response of Cardiac Fibroblasts to Thyroid Hormone in Vitro. Circ Res (1992) 71(4):831–9. Epub 1992/10/01. doi: 10.1161/01.res.71.4.831.

25. Kerp H, Hones GS, Tolstik E, Hones-Wendland J, Gassen J, Moeller LC, et al. Protective Effects of Thyroid Hormone Deprivation on Progression of Maladaptive Cardiac Hypertrophy and Heart Failure. Front Cardiovasc Med (2021) 8:683522. Epub 2021/08/17. doi: 10.3389/fcvm.2021.683522.

26. Lorenz K, Schmitt JP, Schmitteckert EM, Lohse MJ. A New Type of Erk1/2 Autophosphorylation Causes Cardiac Hypertrophy. Nature medicine (2009) 15(1):75–83. Epub 2008/12/09. doi: 10.1038/nm.1893.

27. Schmid E, Neef S, Berlin C, Tomasovic A, Kahlert K, Nordbeck P, et al. Cardiac Rkip Induces a Beneficial Beta-Adrenoceptor-Dependent Positive Inotropy. Nature medicine (2015) 21(11):1298–306. Epub 2015/10/20. doi: 10.1038/nm.3972.

28. Engels K, Rakov H, Hones GS, Brix K, Kohrle J, Zwanziger D, et al. Aging Alters Phenotypic Traits of Thyroid Dysfunction in Male Mice with Divergent Effects on Complex Systems but Preserved Thyroid Hormone Action in Target Organs. The journals of gerontology Series A, Biological sciences and medical sciences (2019). Epub 2019/02/17. doi: 10.1093/gerona/glz040.

29. Rakov H, De Angelis M, Renko K, Hoenes S, Zwanziger D, Moeller LC, et al. Aging Is Associated with Low Thyroid State and Organ Specific Sensitivity to Thyroxine. Thyroid (2019). Epub 2019/08/24. doi: 10.1089/thy.2018.0377.

30. Pantos C, Dritsas A, Mourouzis I, Dimopoulos A, Karatasakis G, Athanassopoulos G, et al. Thyroid Hormone Is a Critical Determinant of Myocardial Performance in Patients with Heart Failure: Potential Therapeutic Implications. Eur J Endocrinol (2007) 157(4):515–20. Epub 2007/09/26. doi: 10.1530/EJE-07-0318.

31. Tomasovic A, Brand T, Schanbacher C, Kramer S, Hummert MW, Godoy P, et al. Interference with Erk-Dimerization at the Nucleocytosolic Interface Targets Pathological Erk1/2 Signaling without Cardiotoxic Side-Effects. Nat Commun (2020) 11(1):1733. Epub 2020/04/09. doi: 10.1038/s41467-020-15505-4.

32. Leask A. Getting to the Heart of the Matter: New Insights into Cardiac Fibrosis. Circ Res (2015) 116(7):1269–76. Epub 2015/03/31. doi: 10.1161/CIRCRESAHA.116.305381.

33. Maisel A. B-Type Natriuretic Peptide Levels: A Potential Novel “White Count” for Congestive Heart Failure. J Card Fail (2001) 7(2):183–93. doi: 10.1054/jcaf.2001.24609.

34. Brandt RR, Wright RS, Redfield MM, Burnett JC, Jr. Atrial Natriuretic Peptide in Heart Failure. J Am Coll Cardiol (1993) 22(4 Suppl A):86A–92A. Epub 1993/10/01. doi: 10.1016/0735-1097(93)90468-g.

35. Lips DJ, deWindt LJ, van Kraaij DJ, Doevendans PA. Molecular Determinants of Myocardial Hypertrophy and Failure: Alternative Pathways for Beneficial and Maladaptive Hypertrophy. European heart journal (2003) 24(10):883–96. doi: 10.1016/s0195-668x(02)00829-1.

36. Visser WE, Bombardieri CR, Zevenbergen C, Barnhoorn S, Ottaviani A, van der Pluijm I, et al. Tissue-Specific Suppression of Thyroid Hormone Signaling in Various Mouse Models of Aging. PLoS One (2016) 11(3):e0149941. Epub 20160308. doi: 10.1371/journal.pone.0149941.

37. Isoyama S, Sato F, Takishima T. Effect of Age on Coronary Circulation after Imposition of Pressure-Overload in Rats. Hypertension (1991) 17(3):369–77. Epub 1991/03/01. doi: 10.1161/01.hyp.17.3.369.

38. Isoyama S, Wei JY, Izumo S, Fort P, Schoen FJ, Grossman W. Effect of Age on the Development of Cardiac Hypertrophy Produced by Aortic Constriction in the Rat. Circ Res (1987) 61(3):337–45. Epub 1987/09/01. doi: 10.1161/01.res.61.3.337.

39. Isoyama S, Nitta-Komatsubara Y. Acute and Chronic Adaptation to Hemodynamic Overload and Ischemia in the Aged Heart. Heart Fail Rev (2002) 7(1):63–9. Epub 2002/01/16.

40. Sopko NA, Turturice BA, Becker ME, Brown CR, Dong F, Popovic ZB, Penn MS. Bone Marrow Support of the Heart in Pressure Overload Is Lost with Aging. PLoS One (2010) 5(12):e15187. Epub 20101221. doi: 10.1371/journal.pone.0015187.

41. Geng X, Hwang J, Ye J, Shih H, Coulter B, Naudin C, et al. Aging Is Protective against Pressure Overload Cardiomyopathy Via Adaptive Extracellular Matrix Remodeling. Am J Cardiovasc Dis (2017) 7(3):72–82. Epub 2017/07/12.

42. Florini JR, Saito Y, Manowitz EJ. Effect of Age on Thyroxine-Induced Cardiac Hypertrophy in Mice. J Gerontol (1973) 28(3):293–7. Epub 1973/07/01. doi: 10.1093/geronj/28.3.293.

43. Bachman ES, Hampton TG, Dhillon H, Amende I, Wang J, Morgan JP, Hollenberg AN. The Metabolic and Cardiovascular Effects of Hyperthyroidism Are Largely Independent of Beta-Adrenergic Stimulation. Endocrinology (2004) 145(6):2767–74. Epub 2004/03/16. doi: 10.1210/en.2003-1670.

44. Herrmann B, Harder L, Oelkrug R, Chen J, Gachkar S, Nock S, et al. Central Hypothyroidism Impairs Heart Rate Stability and Prevents Thyroid Hormone-Induced Cardiac Hypertrophy and Pyrexia. Thyroid (2020) 30(8):1205–16. Epub 20200424. doi: 10.1089/thy.2019.0705.

45. Kerp H, Engels K, Kramer F, Doycheva D, Sebastian Hönes G, Zwanziger D, et al. Age Effect on Thyroid Hormone Brain Response in Male Mice. Endocrine (2019) 66(3):596–606. Epub 20190907. doi: 10.1007/s12020-019-02078-6.

46. Wang X, Wang H, Li Q, Wang P, Xing Y, Zhang F, et al. Effect of Levothyroxine Supplementation on the Cardiac Morphology and Function in Patients with Subclinical Hypothyroidism: A Systematic Review and Meta-Analysis. J Clin Endocrinol Metab (2022) 107(9):2674–83. doi: 10.1210/clinem/dgac417.

47. Cappola AR, Fried LP, Arnold AM, Danese MD, Kuller LH, Burke GL, et al. Thyroid Status, Cardiovascular Risk, and Mortality in Older Adults. Jama (2006) 295(9):1033–41. doi: 10.1001/jama.295.9.1033.

48. Cappola AR, Desai AS, Medici M, Cooper LS, Egan D, Sopko G, et al. Thyroid and Cardiovascular Disease Research Agenda for Enhancing Knowledge, Prevention, and Treatment. Circulation (2019) 139(25):2892–909. Epub 2019/05/14. doi: 10.1161/CIRCULATIONAHA.118.036859.

49. Hönes GS, Härting N, Mittag J, Kaiser FJ. Trα2—an Untuned Second Fiddle or Fine-Tuning Thyroid Hormone Action? International journal of molecular sciences (2022) 23(13):6998.

50. Sylvén C, Jansson E, Sotonyi P, Waagstein F, Barkhem T, Brönnegård M. Cardiac Nuclear Hormone Receptor Mrna in Heart Failure in Man. Life sciences (1996) 59(22):1917–22. Epub 1996/01/01. doi: 10.1016/s0024-3205(96)00539-5.

51. Kinugawa K, Minobe WA, Wood WM, Ridgway EC, Baxter JD, Ribeiro RC, et al. Signaling Pathways Responsible for Fetal Gene Induction in the Failing Human Heart: Evidence for Altered Thyroid Hormone Receptor Gene Expression. Circulation (2001) 103(8):1089–94. Epub 2001/02/27. doi: 10.1161/01.cir.103.8.1089.

52. Kinugawa K, Jeong MY, Bristow MR, Long CS. Thyroid Hormone Induces Cardiac Myocyte Hypertrophy in a Thyroid Hormone Receptor Alpha1-Specific Manner That Requires Tak1 and P38 Mitogen-Activated Protein Kinase. Mol Endocrinol (2005) 19(6):1618–28. Epub 2005/04/16. doi: 10.1210/me.2004-0503.

