## Supplemental Material for "Cardiac recovery from pressure overload is not altered by thyroid hormone status in old mice"

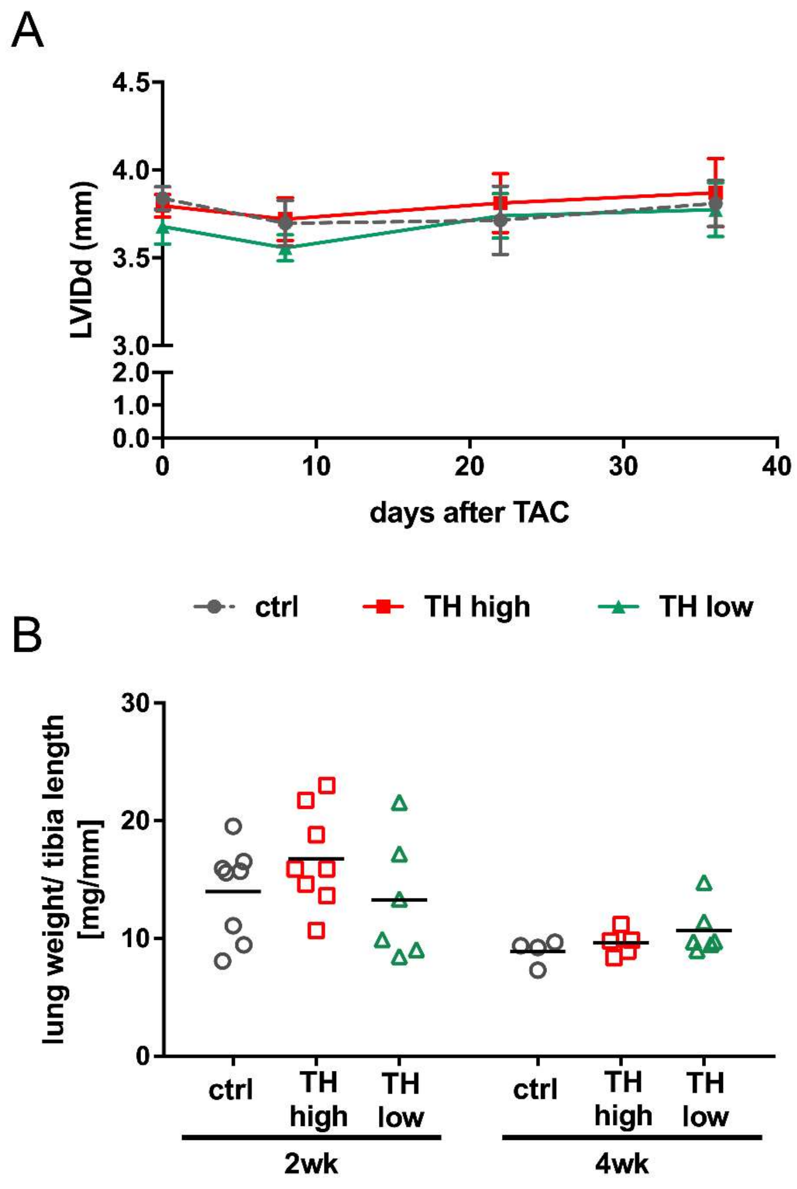


**Suppl. Figure 1:** Left ventricular inner diameter and lung weights of mice subjected to TAC. Diastolic left ventricular inner diameter (LVIDd; A) was quantified by echocardiography during experimental time and lung weight to tibia length ratio at day of sacrifice (B). Values are indicated as mean ± SEM or scatter dot plot and mean with no significant changes according to Two-Way ANOVA and Tukey´s *post hoc* analysis, ctrl=control, wk=weeks.

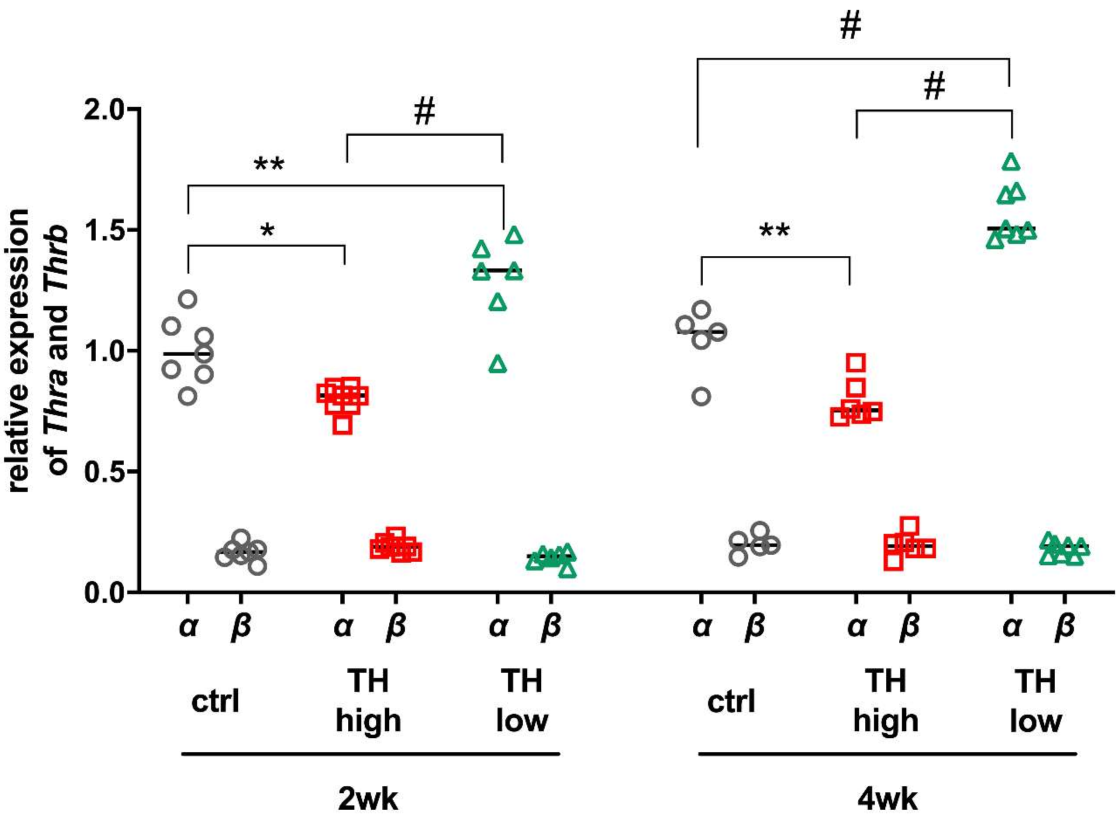


**Suppl. Figure 2:** Expression analysis of TRα and TRβ. Transcripts of total *Thra* and *Thrb* were determined in mouse hearts by qRT-PCR after 2 and 4 weeks of treatment. Scatter dot plot and mean in all panels, One-Way ANOVA with Tukey`s *post-hoc* test, *$p$<0.05, **$p$<0.01, #$p$<0.0001; ctrl=control, wk=weeks.

**Suppl. Table 1:** Oligonucleotides for quantitative RT-PCR. Oligonucleotides were designed using PrimerBlast (NCBI) and synthesized by Eurofins (Eurofins MWG Synthesis, Ebersberg, Germany).

| Gene name | Forward primer (5'-3') | Reverse primer (5'-3') |
|---|---|---|
| *Gapdh* | CCTCGTCCCGTAGACAAAATG | TGAAGGGGTCGTTGATGGC |
| *Rn18S* | CGGCTACCACATCCAAGGAA | GCTGGAATTACCGCGGCT |
| *Polr2a* | CTTTGAGGAAACGGTGGATGTC | TCCCTTCATCGGGTCACTCT |
| *Anp* | TCGTCTTGGCCTTTTGGCTT | GGTGGTCTAGCAGGTTCTTGAAAT |
| *Bnp* | GTTTGGGCTGTAACGCACT | TCACTTCAAAGGTGGTCCCAG |
| *Myh6* | CAGACAGAGATTTCTCCAACCCA | GCCTCTAGGCGTTCCTTCTC |
| *Myh7* | CACGTTTGAGAATCCAAGGCTC | CTCCTTCTCAGACTTCCGCA |
| *Serca2a2* | AACTACCTGGAACAACCCGC | TCATGCAGAGGGCTGGTAGA |
| *Pln* | TTCATGCTCTGCACTGTGACG | GCCAAATGTGAGCTGTCTTCTTTT |
| *Ryr2* | AGGGCAATGAACACTACGGG | CATCCTGCCTACTTGTGCCA |
| *Thra (total)* | GAAAAGCAGCATGTCAGGGTA | GGATTGTGCGGCCAAAGAAG |
| *Thra1* | GCTTCTGACGCCATCTTTGAAC | TCGACTTTCATGTGGAGGAAGC |
| *Thra2* | GCTTCTGACGCCATCTTTGAAC | ATTCCGAGAAGCTGCTGTCC |
| *Thrb* | GGACAAGCACCCATCGTGAA | ACATGGCAGCTCACAAAACAT |

**Suppl. Table 2:** Echocardiographic parameters of all time points. Values are represented as mean ± standard deviation, n=number of animals, nd=not determined. *p<0.05 compared to basal values, #p<0.05 compared to control group at the indicated time (3 or 5 weeks), by One-Way ANOVA and Tukey´s *post hoc* analysis.

| 12 months | basal (n=56) | 1w TAC (n=46) | 3w TAC control (n=13) | 3w TAC TH high (n=15) | 3w TAC TH low (n=15) | 5w TAC control (n=5) | 5w TAC TH high (n=7) | 5w TAC TH low (n=7) |
|---|---|---|---|---|---|---|---|---|
| heart rate [bpm] | 542±65 | 594±78 * | 564±87 | 596±83 | 589±52 | 596±109 | 594±80 | 527±40 |
| IVSd [mm] | 0.70±0.05 | 0.98±0.08 * | 1.01±0.08 * | 1.11±0.12 *,# | 0.97±0.10 * | 1.00±0.07 * | 1.02±0.10 * | 0.88±0.08 * |
| IVSs [mm] | 1.04±0.10 | 1.25±0.12 * | 1.31±0.12 * | 1.39±0.18 * | 1.27±0.13 * | 1.29±0.06 * | 1.26±0.12 * | 1.24±0.18 * |
| LVIDd [mm] | 3.77±0.29 | 3.61±0.43 | 3.71±0.67 | 3.81±0.63 | 3.74±0.48 | 3.81±0.26 | 3.87±0.48 | 3.77±0.38 |
| LVIDs [mm] | 2.46±0.28 | 2.62±0.47 | 2.72±0.76 | 2.72±0.80 | 2.68±0.62 | 2.91±0.42 | 2.87±0.54 | 2.75±0.32 |
| LVPWd [mm] | 0.71±0.04 | 0.98±0.08 * | 1.01±0.10 * | 1.09±0.10 * | 0.95±0.08 * | 0.99±0.08 * | 1.02±0.09 * | 0.85±0.08 *,# |
| LVPWs [mm] | 1.13±0.09 | 1.29±0.11 * | 1.38±0.14 * | 1.44±0.18 * | 1.29±0.12 * | 1.29±0.09 | 1.37±0.13 * | 1.17±0.09 |
| FS [%] | 34.87±4.41 | 28.03±5.57 * | 27.83±7.48 * | 29.99±9.24 | 29.15±7.83 * | 23.97±6.82 * | 26.22±6.49 * | 27.09±4.94 * |
| LV Mass [mg] | 90.49±11.19 | 133.41±27.59 * | 144.89±31.33 * | 171.48±35.47 * | 136.03±20.71 * | 147.67±16.84 * | 157.89±34.38 * | 118.60±12.40 |
| LV Mass corrected [mg] | 72.39±8.95 | 106.73±22.07 * | 115.91±25.06 * | 137.19±28.38 * | 108.82±16.57 * | 118.13±13.47 * | 126.31±27.51 * | 94.88±9.92 |
| LV Vol d [µl] | 61.24±11.36 | 56.10±16.43 | 61.64±31.27 | 65.02±26.86 | 61.13±19.82 | 62.78±10.07 | 66.23±19.27 | 61.88±13.11 |
| LV Vol s [µl] | 21.84±6.22 | 26.35±12.07 | 31.10±25.22 | 31.53±23.52 | 28.90±18.09 | 33.59±11.33 | 33.28±16.21 | 28.91±7.31 |
| AV Peak Velocity [mm/s] | nd | -4539±353 | -4533±297 | -4366±573 | -4806±387 | -4475±532 | -4438±241 | -4420±271 |
| AV Peak Pressure [mmHg] | nd | 82.91±12.98 | 82.53±11.10 | 77.25±17.10 | 92.99±14.67 | 81.23±19.29 | 79.01±8.68 | 78.44±9.82 |
